# Atg8a interacts with transcription factor Sequoia to control the expression of autophagy genes in *Drosophila*

**DOI:** 10.1101/611731

**Authors:** Anne-Claire Jacomin, Stavroula Petridi, Marisa DiMonaco, Ashish Jain, Zambarlal Bhujabal, Nitha Mulakkal, Anthimi Palara, Emma L. Powell, Bonita Chung, Cleidiane G. Zampronio, Alexandra Jones, Alexander Cameron, Terje Johansen, Ioannis P. Nezis

**Affiliations:** School of Life Sciences, University of Warwick, CV4 7AL Coventry, United Kingdom; Molecular Cancer Research Group, Institute of Medical Biology, University of Tromsø – The Arctic University of Norway, 9037 Tromsø, Norway; Department of Molecular Cell Biology, Institute for Cancer Research, Oslo University Hospital, Montebello, N-0379 Oslo, Norway; Centre for Cancer Biomedicine, Faculty of Medicine, University of Oslo, Montebello, N-0379 Oslo, Norway

**Author notes:** These authors contributed equally to this work (Anne-Claire Jacomin, Stavroula Petridi).

**Keywords:** acetylation, autophagy, LIR motif, nucleus, transcription, LC3/Atg8

## Abstract

Autophagy is a fundamental, evolutionarily conserved, process in which cytoplasmic material is degraded through the lysosomal pathway [1–7]. One of the most important and well-studied autophagy-related proteins is LC3 [Microtubule-associated protein 1 light <u>c</u>hain <u>3,</u> (called Atg8 in yeast and *Drosophila*)], which participates in autophagosome formation and autophagy cargo selection in the cytoplasm, and is one of the most widely utilized markers of autophagy [8, 9]. Despite growing evidence that LC3 is enriched in the nucleus, little is known about the mechanisms involved in targeting LC3 to the nucleus and the nuclear components it interacts with [10–13]. Here we show that *Drosophila* Atg8a protein, homologous to mammalian LC3 and yeast Atg8, interacts with the transcription factor Sequoia in a LIR-motif dependent manner. We show that Sequoia depletion induces autophagy in nutrient rich conditions through enhanced expression of autophagy genes. We also show that Atg8a interacts with YL-1, a component of a nuclear acetyltransferase complex, and is acetylated at position K46. Additionally, we show that Atg8a interacts with the deacetylase Sir2, which deacetylates Atg8a during starvation in order to activate autophagy. Our results suggest a mechanism of regulation of expression of autophagy genes by Atg8a, which is linked to its acetylation status and its interaction with Sequoia, YL-1 and Sir2.

## RESULTS AND DISCUSSION

### Sequoia is an Atg8a-interacting protein

In order to identify novel Atg8a-interacting proteins in *Drosophila*, we screened the *Drosophila* proteome for LIR motif-containing proteins using the iLIR software that we have previously developed [14, 15]. We found that the transcription factor ZnF-C2H2-containing protein Sequoia (CG32904) has a predicted LIR motif at position 311-316 with the sequence EEYQVI (Fig. S1A) [14–16]. Sequoia contains two zinc-finger domains which are homologous to the DNA-binding domain of Tramtrack and has been shown to regulate neuronal morphogenesis [17]. We confirmed the direct interaction between Sequoia and Atg8a using GST pulldown binding assays (Fig. 1A, B). This interaction was greatly reduced when we used a form of Atg8a where the LIR docking site (Y49A) was impaired, indicating that the interaction between Sequoia and Atg8a is LIR motif-dependent [18–20]. Futhermore, point mutations of Sequoia LIR motif in positions 313 and 316 by alanine substitutions of the aromatic- and hydrophobic residues (Y313A and I316A) reduced its binding to Atg8a (Fig. 1A, B).

**Figure 1.**
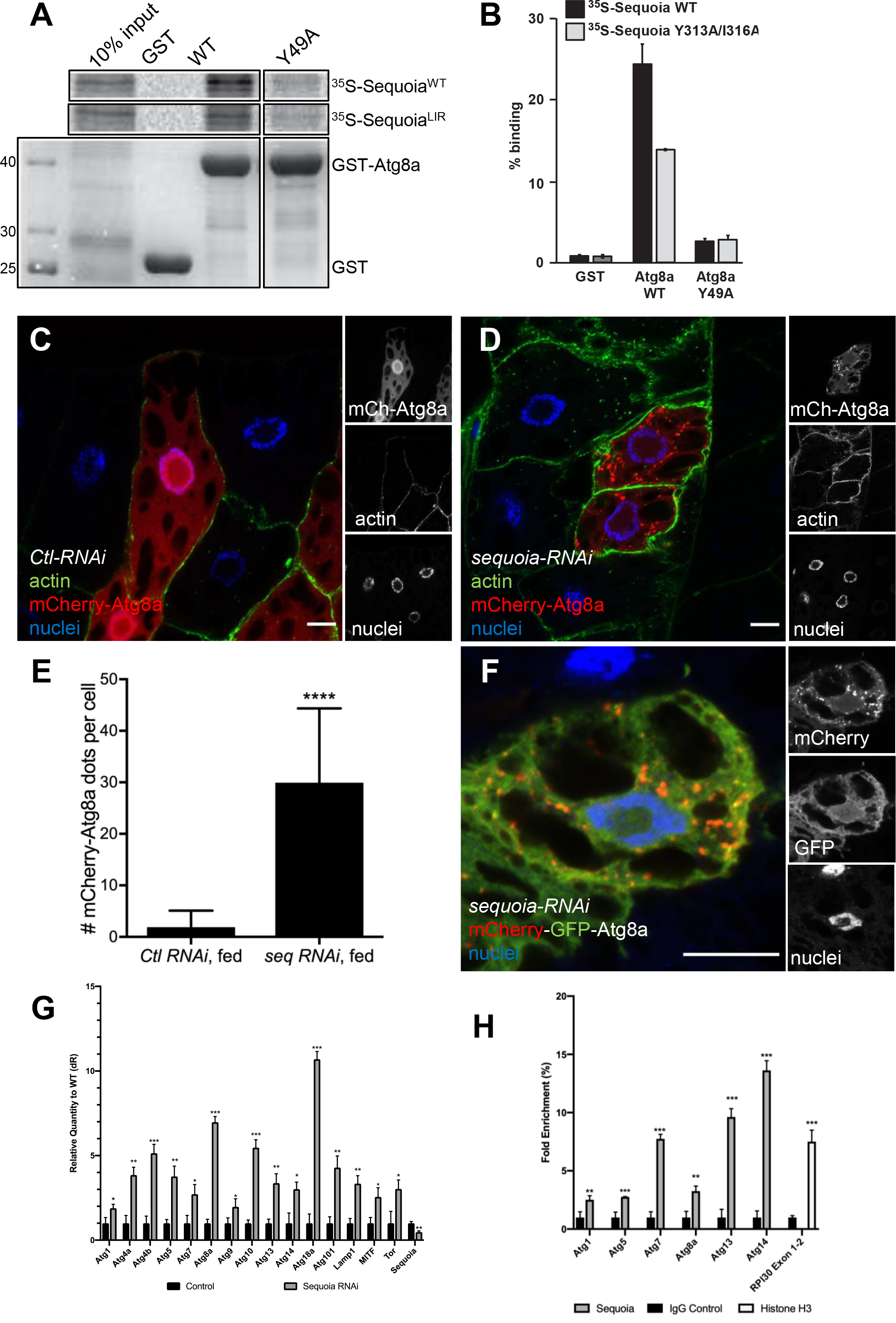
Sequoia negatively regulates autophagy. (A-B) Sequoia interacts with Atg8a in a LIR-dependent manner. (A) GST-pull-down assay between GST-tagged Atg8a-WT or -LDS mutant (Y49A), and radiolabelled myc-Sequoia-WT or -LIR mutant (Y313A/316A). GST was used as negative control. Quantification of the binding is shown in (B). (C-D) Confocal sections of larval fat bodies clonally expressing the autophagy marker mCherry-Atg8a (red) in combination with a control RNAi (C) or a sequoia RNAi (D). Fixed fat bodies where stained for cortical actin (green) and nuclei (blue). Scale bar: 10µm. (E) Quantification of the number of mCherry-Atg8a dots per cell. Bars denote mean ± s.d. Statistical significance was determined using Student’s t-test: ****p < 0.0001. (F) Confocal sections of larval fat bodies clonally expressing the autophagy flux marker GFP-mCherry-Atg8a (red and green) in combination with a sequoia RNAi. Fixed fat bodies where stained for nuclei (blue). Scale bar: 10µm. (G) Analysis of the mRNA level of autophagy-associated genes in control and sequoia-depleted fat bodies using real time qPCR. (H) Analyses of Sequoia binding to the promoter of autophagy genes as detected by ChIP (chromatin immunoprecipitation). IgG was used as a negative control and Histone H3 binding to RPL30 was used as a positive control. Genotypes: (C) *y w hs-Flp; Ac>CD2>GAL4 / +; UAS-mCherry-Atg8a / UAS-luc-RNAi*, (D) *y w hs-Flp; Ac>CD2>GAL4 / +; UAS-mCherry-Atg8a / UAS-sequoia-RNAi*, (F) *y w hs-Flp; Ac>CD2>GAL4 / +; UAS-mCherry-GFP-Atg8a / UAS-sequoia-RNAi*, (G, H) ctl: *Cg-GAL4 /+; UAS-luc-RNAi /+*, seq-RNAi: *Cg-GAL4* /+; UAS-sequoia-RNAi /+

Given the observed interaction between Sequoia and Atg8a, we examined whether Sequoia is degraded by autophagy. Western blot analysis showed that endogenous Sequoia is not accumulated in *Atg8a* and *Atg7* mutants compared to wild-type flies (Fig. S1B). Taken together these results indicate that Sequoia is an Atg8a-interacting protein and that this interaction is LIR-motif-dependent. Interestingly, in spite of its interaction with Atg8a, Sequoia is not a substrate for autophagic degradation.

### Sequoia is a negative transcriptional regulator of autophagy

In order to examine the role of Sequoia in autophagy, we silenced *sequoia* using RNAi alongside with the expression of the autophagic marker mCherry-Atg8a [21, 22]. We observed a significant increase of mCherry-Atg8a puncta in Sequoia-depleted fat body cells in fed conditions compared to control cells (Fig. 1C-E). An accumulation of Atg8a-positive puncta in the cell can result either from an induction or blockade of the autophagic flux. To make the distinction between these two possibilities, we made use of a tandem-tagged Atg8a (GFP-mCherry-Atg8a) [23]. In cells expressing RNAi against Sequoia, we noticed an accumulation of red puncta that lack GFP fluorescence, suggesting an induction of autophagic flux (Fig. 1F). These results show that Sequoia is a negative regulator of autophagy.

We next examined whether expression of autophagy genes was affected upon *Sequoia* knockdown. Real-time quantitative PCR analysis showed that the expression of numerous autophagy genes was increased when *sequoia* was silenced (Fig. 1G). As Sequoia is a transcription factor and contains C2H2 zinc-finger domains putatively involved in DNA-binding, we performed a ChIP assay to test the ability of Sequoia to bind the promotor region of autophagy genes. We found that Sequoia is enriched on the promoters of several autophagy genes suggesting that Sequoia is acting as a repressor to negatively regulate autophagy (Fig. 1H).

### The repressive activity of Sequoia on autophagy depends on its LIR motif

In order to evaluate whether the role of Sequoia in the negative transcriptional regulation of autophagy depends on its interaction with Atg8a via its LIR motif, we created transgenic flies allowing for the expression of GFP-tagged wild-type (GFP-Sequoia-WT) or LIR mutant (GFP-Sequoia-Y313A/I316A) Sequoia under the control of a UAS region. These transgenes were expressed concomitantly with the autophagic marker mCherry-Atg8a. We bserved that both GFP-Sequoia-WT and -Y313A/I316A proteins localized exclusively in the nucleus of fat body cells (Figs. 2A-B). Interestingly, the expression of GFP-Sequoia-Y313A/I316A resulted in a significant increase of mCherry-Atg8a puncta in fed conditions while the expression of wild-type Sequoia had no effect (Figs. 2A-D). Additionally, cells expressing LIR-mutated Sequoia showed an increase of the lysosomal markers Cathepsin L and LysoTracker Red. (Figs. 2E-G). Collectively, these results suggest that Sequoia negatively regulates autophagy through its LIR-motif-dependent interaction with Atg8a.

**Figure 2.**
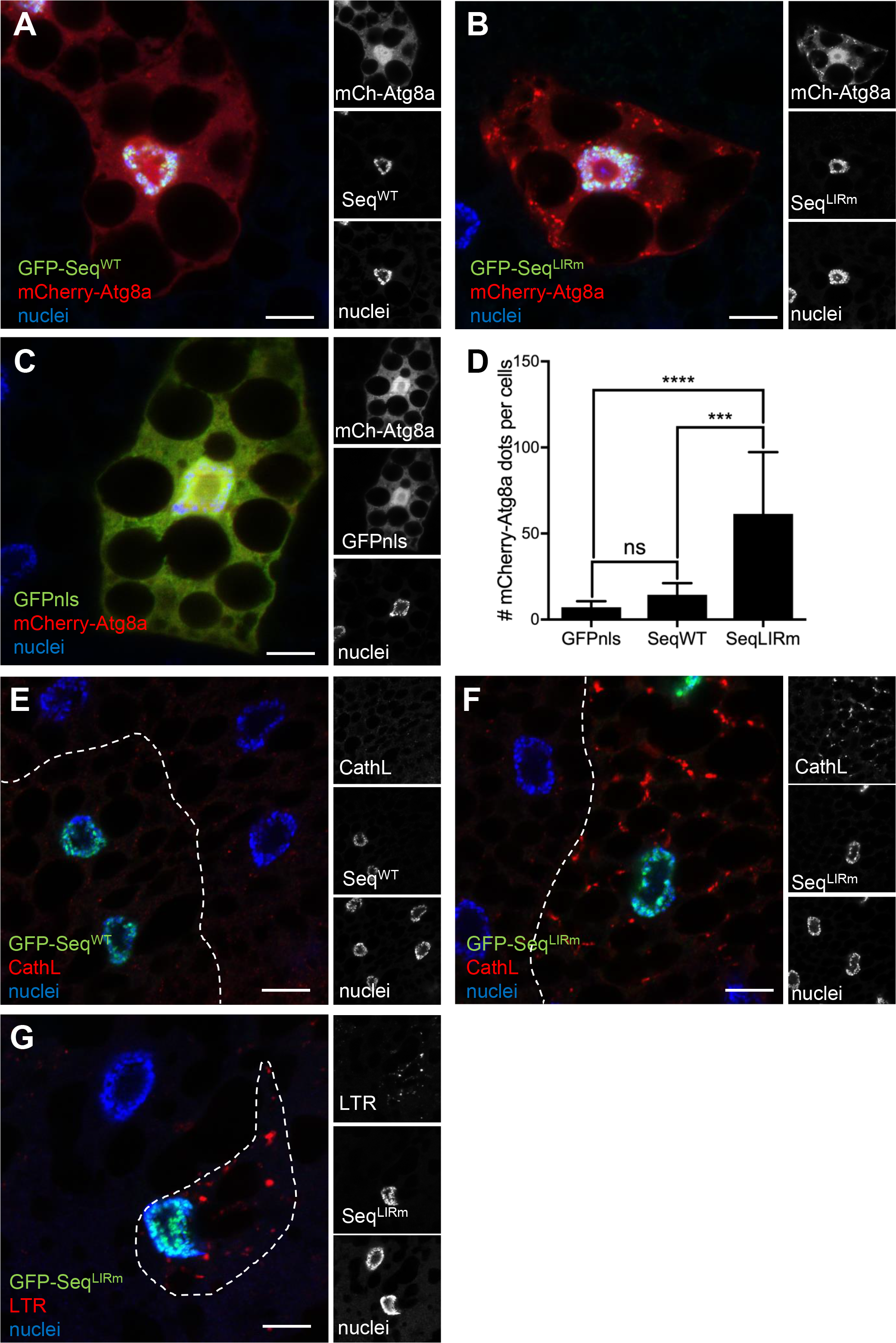
Sequoia’s LIR motif is required to repress autophagy. (A-C) Confocal sections of larval fat bodies clonally expressing the autophagy marker mCherry-Atg8a (red) in combination with GFP-Sequoia WT (A), GFP-Sequoia LIRm (B) or GFPnls (C) (green). Fixed fat bodies where stained for nuclei (blue). Scale bar: 10µm. (D) Quantification of the number of mCherry-Atg8a dots per cell. Bars denote mean ± s.d. Statistical significance was determined using one-way ANOVA: ***p < 0.001, ****p < 0.0001. (E-G) Confocal sections of larval fat bodies clonally expressing GFP-Sequoia WT (E), GFP-Sequoia LIRm (F) (green) and stained for cathepsin-L (red) (E-F, CathL) or LysoTracker Red (G, LTR). Genotypes: (A) *y w hs-Flp; Ac>CD2>GAL4 / UAS-GFP-Sequoia-WT; UAS-mCherry-Atg8a /+*, (B) *y w hs-Flp; Ac>CD2>GAL4 / UAS-GFP-Sequoia-LIRm; UAS-mCherry-Atg8a /+*, (C) *y w hs-Flp; Ac>CD2>GAL4 / +; UAS-mCherry-Atg8a / UAS-GFPnls*, (E) *y w hs-Flp; Ac>CD2>GAL4 / UAS-GFP-Sequoia-WT*, (F, G) *y w hs-Flp; Ac>CD2>GAL4 / UAS-GFP-Sequoia-LIRm*

### Atg8a interacts with the nuclear acetyltransferase subunit YL-1 and is acetylated at K46 in nutrient rich conditions

In a search for other nuclear interactors of Atg8a, we identified the nuclear acetyltransferase subunit YL-1. YL-1 belongs to the multi-subunit chromatin-remodeling complexes called SWR1 in yeast and the related SRCAP and NuA4/Tip60 complexes in mammals that has been shown to control histone acetylation [24–28]. *In vitro* translated YL-1 (^35^S-Myc-YL1) bound very strongly and directly to the N-terminal half (amino acid residues 1-71) of recombinant GST-Atg8a (Fig. 3A, B). However, mutation of either the LDS of Atg8a (Fig. 3A), or the putative LIR motifs of YL-1 (Fig. S2) did not abrogate the *in vitro* interaction between Atg8a and YL-1, suggesting that this interaction is LIR-motif independent.

**Figure 3.**
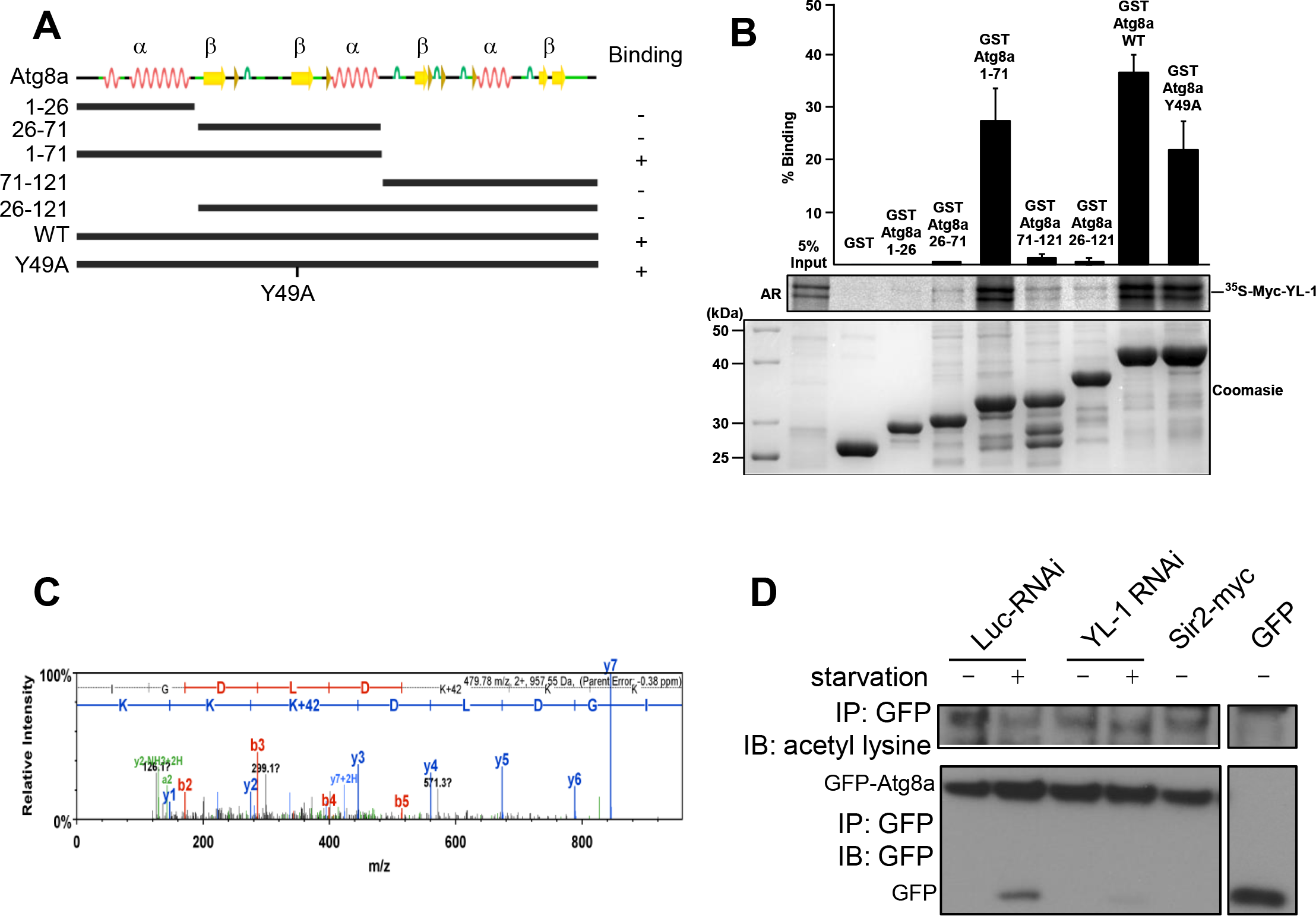
YL-1 interacts with Atg8a. (A) Sequence view of DmAtg8a based on GABARAP structure (PDB:1KOT) with deletion constructs used in GST-pull down assay with YL-1. (B) GST-pull-down assay between GST-tagged Atg8a-WT, deletion constructs or -LDS mutant (Y49A), and radiolabelled myc-YL-1 WT. (C) Acetylation of Atg8a was identified at K46. Fed larvae that were constitutively expressing GFP-Atg8a in their fat bodies were collected and protein acetylation was determined by immunoprecipitation (IP) with a GFP antibody followed by mass spectrometry analysis. Residue K+42, which is the K46 lysine, was identified acetylated. (D) Knockdown of YL-1 reduces acetylation of Atg8a. Flies that were constitutively expressing GFP-Atg8a in their fat bodies were crossed with the control RNAi, the YL-1 RNAi line and the Sir2-Myc line and their offspring were collected. Protein acetylation was determined by immunoprecipitation (IP) with a GFP antibody followed by Western blotting (WB) using an antibody recognizing acetyl-lysine residues. Immunoprecipitation was performed with 1 mg of protein lysate from whole larvae both in fed and starved (4h in 20% sucrose) conditions. Genotypes: (C) *Cg-GAL4 UAS-GFP-Atg8a/+* (D) luc-RNAi: *Cg-GAL4 UAS-GFP-Atg8a/+;UAS-luc-RNAi /+*, YL-1-RNAi: *Cg-GAL4 UAS-GFP-Atg8a/+;UAS-yl-1-RNAi /+*,Sir2-myc: *Cg-GAL4 UAS-GFP-Atg8a/UAS-Sir2-myc*

Since YL-1 is a component of the NuA4/Tip60 acetyltransferase complex, we examined whether Atg8a is acetylated. Mass spectrometry analysis of immunoprecipitated GFP-Atg8a from fed larvae revealed that Atg8a is acetylated on the lysine at position K46 in nutrient rich conditions (Fig. 3C).

To examine whether YL-1 regulates the acetylation of Atg8a, we immunoprecipitated GFP-Atg8a and used an anti-acetyl-lysine antibody to reveal Atg8a acetylation by western blotting. We found that GFP-Atg8a is acetylated in fed conditions and that its acetylation is reduced after starvation and when YL-1 is depleted using RNAi against YL-1 (Fig 3D). All together these data show that YL-1 interacts with and regulates acetylation of Atg8a.

### Sir2 interacts with and deacetylates Atg8a during starvation

To further investigate the impact of acetylated Atg8a on the activation of autophagy, we focused on the deacetylase Sir2, homologue of mammalian Sirtuin-1, that has been shown to have a role in lipid metabolism and insulin resistance [29–31]. Using a fly line for the expression of myc-tagged Sir2 (Sir2-myc), we found that Sir2 localises in the nucleus of fat body cells in both fed and starved larvae (Fig. S3A, B). Fat bodies clonally overexpressing Sir2-myc and stained for acetylated-lysine revealed a reduction of nuclear staining for acetylated protein in clonal cells compared to their wild-type neighbours (Fig. S3C-E). This suggests that dSir2 is required for deacetylation in the nucleus.

Staining fat bodies from starved larvae with LysoTracker Red failed to show an accumulation of lysosomes in the fat bodies from *Sir2* mutant larvae (Fig 4A-C). This observation shows that Sir2 is required for the activation of starvation-induced autophagy.

**Figure 4.**
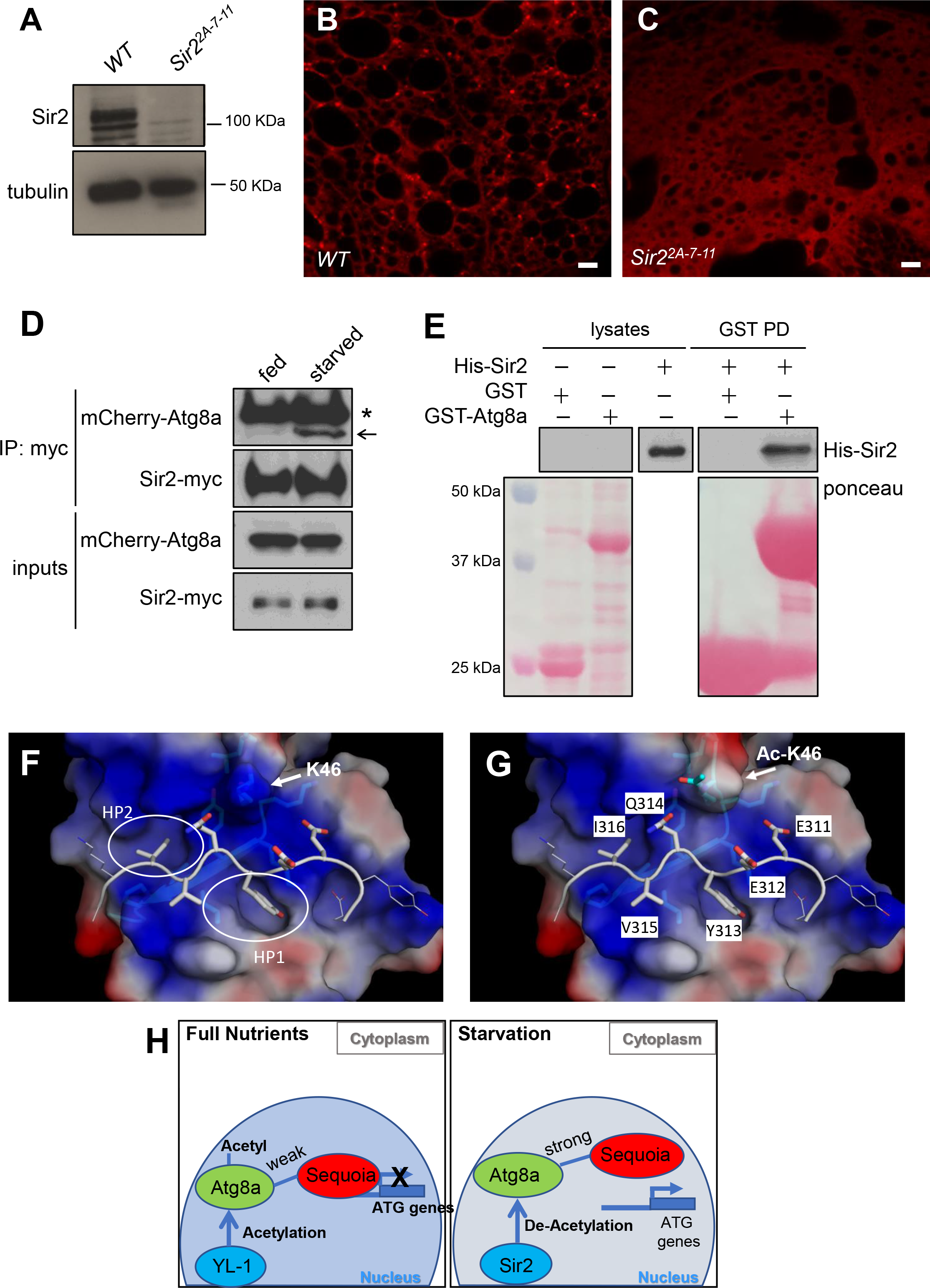
Sir2 interacts with Atg8a and regulates autophagy induction. (A) Sir2 western blotting on protein lysates prepared from 7-day old w1118 and the Sir2[2A-7-11] mutant. Tubulin was used as a loading control. (B-C) Confocal sections of fat bodies from *w*^*1118*^ (b) and the *Sir2*^*2A-7-11*^ (c) larvae stained with LysoTracker Red after 4h starvation. Scale bar: 10µm (D) Co-immunoprecipitation of Sir2-myc and mCherry-Atg8a from extracts of fed and starved larvae. Sir2-myc was immunoprecipitated (IP) from larva lysates and subjected to SDS-PAGE. *: non specific (IgG), arrow: mCherry-Atg8a. (E) *In vitro* interaction between recombinant GST-Atg8a and 6×His-Sir2. GST or GST-Atg8a were pulled down and incubated with bacterial lysate expressing 6×His-Sir2. Eluates were subjected to SDS-PAGE. Membrane was stained with Ponceau S for total protein staining, and immunoblotted with anti-His antibody for His-Sir2. (F-G) Homology model of a peptide of Sequoia binding to Drosophila Atg8a LIR complex in fed (F) and starved (G) conditions. Atg8a is represented by a semi-transparent surface coloured according to electrostatic charge; where blue is positive, white is neutral and red is negative. The grey peptide back-bone structure is representative of the Sequoia LIR peptide. (H) Proposed model of Sequoia/Atg8a/YL-1/Sir2 in induction of autophagy. In fed conditions histone-interacting protein YL-1 recruits and acetylates Atg8a at K46 and Atg8a recruits Sequoia on the promoters of autophagy genes to repress their expression. During starvation, Sir2 interacts with and deacetylates Atg8a. Deacetylated Atg8a is now able to strongly interact with Sequoia, and as it moves to the cytoplasm, it can displace Sequoia from autophagy genes promoters therefore allowing their transcription. Genotypes: (A, B) WT: *w*^*1118*^, (D) *w-;Cg-GAL4 UAS-mCherry-Atg8a/UAS-Sir2-myc*

Additionally, overexpression of Sir2-myc resulted in a reduction of GFP-Atg8a acetylation in fed condition (Fig. 3D). Moreover, we observed that Sir2-myc interacts preferentially with mCherry-Atg8a in starved conditions (Fig 4D). Using *in vitro* GST pulldown assay, we showed that Sir2 interacts directly with Atg8a (Fig. 4E). All together, these results suggest that deacetylation of Atg8a by Sir2 is required for the activation of autophagy.

To examine the significance of K46 acetylation, we created a homology model of the LIR peptide of Sequoia binding to *Drosophila* Atg8a based on the structure of GABARAP-L1 ATG4B LIR Complex (PDB 5LXI) (Fig 4F, G). The residues Y313 and I316 of Sequoia’s LIR peptide are likely to bind in the HP1 and HP2 pockets respectively. The negatively charged glutamates (E) will interact with the positively charged lysine (K) residues of the 44-LDKKKYLVP-52 motif (shown as sticks under the surface). Upon acetylation the K46 residue will become bulkier, will lose the positive charge and this could modulate the binding between Atg8a and Sequoia. These data suggest that deacetylated Atg8a could bind stronger to Sequoia.

Atg8-family proteins have been extensively described for their implication in autophagosome formation and cargo selection in the cytoplasm. Although Atg8 family proteins also localise in the nucleus, their role in this compartment remains largely unexplored.

Here, we uncovered a nuclear role of *Drosophila* Atg8a in the regulation of autophagy genes expression and induction of autophagy via a LIR motif -dependent mechanism, regulated by Atg8a acetylation. We showed that the transcription factor Sequoia interacts with Atg8a in the nucleus in a LIR-dependent manner to repress the transcriptional activation of autophagy genes. We suggest that the acetylation status of Atg8a at position K46 contributes to the modulation of the interaction between Sequoia and Atg8a in the nucleus. We also identified that the nuclear acetyltransferase subunit YL-1 and deacetylase Sir2 interact with Atg8a, and that they act as modulators of Atg8a acetylation. We propose a model (Fig 4H) where in fed conditions histone-interacting protein YL-1 recruits and controls the acetylation of Atg8a at K46 and in turn Atg8a recruits Sequoia on the promoters of autophagy genes to participate in the repression of their expression. During starvation, when autophagy requires to be activated, Sir2 interacts with and deacetylates Atg8a. Deacetylated Atg8a is now able to strongly interact with Sequoia, and as it moves to the cytoplasm, can displace Sequoia from the promotor region of autophagy genes thereby allowing their transcription. Our results suggest a novel nuclear role for Atg8a and a LIR-motif dependent mechanism of regulation of autophagy gene expression, which is linked to the acetylation/deacetylation status of Atg8a and its interaction with transcription repressor Sequoia.

Our results support previous findings about the yeast and mammalian homologues of Sequoia, Rph1 and KDM4A respectively which have been shown to negatively regulate the transcription of autophagy genes [32]. Our study also supports previous reports about the role of acetylation and deacetylation of LC3 in mammals [10, 33–36] and the regulation of autophagy by acetylation [28, 37].

Higher eukaryotes express YL-1, a highly conserved Swc2 homolog, which has specific H2A.Z-binding properties. *Drosophila* YL-1 has been shown to have a H2A.Z-binding domain that binds H2A.Z-H2B dimer [26]. Here we report a novel role for YL-1 in the regulation of acetylation of non-histone proteins and the regulation of autophagy induction.

In conclusion, we have shown a role of Atg8a in regulation of the transcription of autophagy genes in *Drosophila*. Our study highlights the physiological importance of a non-degradative LIR-motif dependent interaction of Atg8a with a transcription factor and provide novel insights on an unanticipated nuclear role of a protein that controls cytoplasmic cellular self-eating.

## ACKNOWLEDGMENTS

This work was supported by BBSRC grant BB/P007856/1 and Leverhulme Trust project grant RPG-2017-023 to I.P.N. We thanks Prof J Yan (University of California) for the Sequoia antibody. We thank Bloomington stock centre and Vienna RNAi stock centre for flies. We thank Mark Ward for preparing fly food.

## AUTHOR CONTRIBUTIONS

ACJ, SP, MD, AJ, BZ, NM, AP, EP, BC, CZ, AJ, TJ and IPN performed experiments and/or analysed data. AC did the structural modelling. IPN designed and supervised the study. ACJ, SP and IPN drafted the manuscript. All authors read and contributed to the manuscript.

## STAR METHODS

### Fly husbandry and generation of transgenic lines

Flies used in experiments were kept at 25 °C and 70% humidity raised on cornmeal-based feed. The following fly stocks were obtained from the Bloomington *Drosophila* stock center: *w*^1118^ (#3605), *Cg-GAL4* (#7011) and *UAS-YL-1-RNAi* (#31938), *UAS-Sir2-myc (#44216)*. *UAS-Sequoia-RNAi* (#50146) was obtained from the Vienna Drosophila Resource Center. The following mutant lines have been used: *Atg7*^*Δ77*^, and *Atg7*^*Δ14*^/*CyO-GFP* [38], *Sir2*^*2A-7-11*^ from Bloomington (#8838). The clonal analysis using the FLPout system has been performed with the following lines: *yw hs-flp; UAS-mCherry-Atg8a;Ac > CD2 > GAL4*. The transgenic lines UAS-GFP-Sequoia-WT, UAS-GFP-Sequoia-LIRm (Y313A/316A) and UAS-GFP-YL-1 have been generated by cloning the cDNA of *sequoia* or *YL-1* respectively into the pPGW plasmid (DGRC). Transgenic flies were generated by P-element-mediated transformation (BestGene Inc). Early third-instar larvae were collected either fed or starved for 4 hours in 20% sucrose solution in PBS.

### Protein extraction, immunoprecipitation, and western blotting

Protein content was extracted from larvae in Nuclear lysis buffer (20 mM Tris pH 7.5, 137 mM NaCl, 1mM MgCl_2_, 1mM CaCl_2_, 1% Igepal, 10% Glycerol, 1mM Na_3_VO_4_, 15mM Na_4_P_2_O_7_, 5mM Sodium butyrate supplemented with EDTA-free proteases inhibitors cocktail (Roche, 5892791001) and deacetylase inhibitors (Santa Cruz, sc-362323) using a motorized mortar and pestle. Co-immunoprecipitations were performed on lysates from flies expressing GFP alone, GFP-Atg8a or Sir2-myc along with mCherry-Atg8a. After a 30 min pre-clear of the lysates (1 mg total proteins) with sepharose-coupled G-beads (Sigma), the co-immunoprecipitations were performed for 2 h at 4 °C using an anti-GFP antibody (Abcam, Ab290) or anti-myc (Cell Signaling Technology, #2276) and fresh sepharose-coupled G-beads. Four consecutive washes with the lysis buffer were performed before suspension of the beads in 60 μL 2X Laemmli loading buffer. All protein samples (whole fly lysates and co-immunoprecipitation eluates) were boiled for 5-10 min at 95 °C. Quantity of 10–40 μg of proteins (whole lysates) or 20 μL (co-immunoprecipitation eluates) were loaded on polyacrylamide gels and were transferred onto either nitrocellulose or PVDF membranes (cold wet transfer in 10-20% ethanol for 1h at 100V). Membranes were blocked in 5% BSA or non-fat milk in TBST (0.1% Tween-20 in TBS) for 1 h. Primary antibodies diluted in TBST were incubated overnight at 4 °C or for 2 h at room temperature with gentle agitation. HRP-coupled secondary antibodies binding was done at room temperature (RT) for 45 min in 1% BSA or non-fat milk dissolved in TBST and ECL mix incubation for 2 min. All washes were performed for 10 min in TBST at RT. The following primary antibodies were used: anti-GFP (Santa Cruz sc-9996, 1:1000), anti-mCherry (Novus NBP1 #96752, 1:1000), anti-alpha tubulin (Sigma-Aldrich T5168, 1:50,000), anti-acetyl lysine (Cell Signalling #9441, 1:1000), anti-dSir2 (DSHB p4A10, 1:50). HRP-coupled secondary antibodies were from Thermo Scientific (anti-mouse HRP #31450; anti-rabbit HRP #31460). Following co-immunoprecipitation, Veriblot HRP-coupled IP secondary antibody was used (Abcam ab131366, 1:5000).

### Immunohistochemistry

Larva tissues were dissected in PBS and fixed for 30 min in 4% formaldehyde in PBS. Blocking and antibody incubations were performed in PBT (0.3% BSA, 0.3% Triton X-100 in PBS). Primary and secondary antibodies were incubated overnight at 4°C in PBT. The following primary antibody was used: anti-Cathepsin-L (Abcam ab58991, 1:400). Washes were performed in PBW (0.1% Tween-20 in PBS). All images were acquired using Carl Zeiss LSM710 or LSM880 confocal microscopes, using a ×63 Apochromat objective. Staining with LysoTracker was performed by incubating the non-fixed larval fat body in LysoTracker Red in PBS (1:1,000) for 10 min followed by mounting in 75 % glycerol in PBS. Fiji/ImageJ was used to quantify the mCherry-Atg8a dots using the macro AtgCOUNTER [39].

### Real-Time qPCR

RNA extraction was performed with a Life Technologies Ambion PureLinkTM RNA Mini kit according to the manufacturer protocol. Fat bodies from 20 L3 larvae were used per extract. Subsequent steps were performed using 1 μg of total RNA. ThermoScientific DNase I was used in order to digest genomic DNA. The ThermoScientific RevertAid Kit was subsequently used to synthesize cDNA. RT-qPCR was performed using the Promega GoTaq qPCR Master Mix (ref. A6002). Primer sequences are available in Supplementary Table 1.

### Plasmid constructs

Sequences of the genes of interest were amplified by PCR and inserted in desired plasmid using either Gateway recombination system or restriction enzyme cloning. PCR products were amplified from cDNA using Phusion high fidelity DNA polymerase with primers containing the Gateway recombination site or restriction enzyme sites for Gateway entry vector and cloned into pDONR221 or pENTR using Gateway recombination cloning. Point-mutants were generated using the QuikChange site-directed mutagenesis (Stratagene, 200523). Plasmid constructs were verified by conventional restriction enzyme digestion and/or by DNA sequencing.

### GST Pull-down assays

GST-fusion proteins were expressed in Escherichia coli BL21(DE3), SoluBL21 or Rosetta2. GST-fusion proteins were purified on glutathione-Sepharose 4 Fast Flow beads (Amersham Biosciences). GST pull-down assays were performed using either recombinant proteins produced in bacteria or *in vitro* translated ^35^S-methionine-labelled proteins. L-[35S]-methionine was obtained from PerkinElmer Life Sciences. A volume of 10 μL of the *in vitro* translation reaction products (0.5 μg of plasmid in a 25 μL reaction volume) were incubated with 1–10 μg of GST-recombinant protein in 200 μL of NETN buffer (50 mM Tris, pH 8.0, 100 mM NaCl, 6 mM EDTA, 6 mM EGTA, 0.5% Nonidet P-40, 1 mM dithiothreitol supplemented with Complete Mini EDTA-free protease inhibitor cocktail (Roche Applied Science)) for 1 h at 4 °C, washed six times with 1 ml of NETN buffer, boiled with 2X SDS gel loading buffer, and subjected to SDS-PAGE. Gels were stained with Coomassie Blue and vacuum-dried. ^35^S-Labeled proteins were detected on a Fujifilm bio-imaging analyzer BAS-5000 (Fuji).

### Identification of proteins and acetylation using LC-MS/MS

The purified GFP-Atg8a from larvae was separated on 8% polyacrylamide gels. The gel bands corresponding to the targeted protein were stained with Coomasie blue and excised from the gel, slices were extensively washed and proteins were reduced with 10 mM of TCEP and alkylated with 40 mM CAA. Then in-gel digestion was carried out using sequence grade modified trypsin (Promega, Fitchburg, WI) in 50 mM ammonium bicarbonate at 37°C overnight. The peptides were extracted twice with 5% formic acid in 25% acetonitrile aqueous solution and volume of the extractions were reduced in a speedvac. For LC-MS/MS analysis, the peptides were separated by a 60 min gradient elution at a flow rate 250 nL/min with a Thermo-Dionex Ultimate 3000 HPLC system, which was directly interfaced with an Orbitrap Fusion mass spectrometer (Thermo Scientific). The sample was loaded on an Acclaim-PepMap µprecolumn 300 µm id × 5mm length, 5 µm particle size, 100 Å pore size (Thermo Scientific) equilibrated in 2% ACN and 0.1% TFA for 8 min at 10 µL/min. The analytical column utilised was an Acclaim PepMap RSLC 75 µm id × 50 cm, 2 µm 100 Å (Thermo Scientific). The columns were installed on an Ultimate 3000-RSLCnano system (Dionex). Mobile phase A consisted of 0.1% formic acid in water, and mobilephase B consisted of 100% acetonitrile and 0.1% formic acid. The Orbitrap Fusion mass spectrometer was operated in the data-dependent acquisition mode using Xcalibur 2.2 software. Survey scans of peptide precursors from 375 to 1500 *m*/*z* were performed at 120K resolution (at 200 *m*/*z*) with a 2 × 10^5^ ion count target. Tandem MS was performed by isolation at 1.2 Th using the quadrupole, HCD fragmentation with normalized collision energy of 33, and rapid scan MS analysis in the ion trap. The MS^2^ ion count target was set to 5×10^3^ and the max injection time was 200 ms. The dynamic exclusion duration was set to 40 s with a 10 ppm tolerance around the selected precursor and its isotopes. Monoisotopic precursor selection was turned on and the instrument was run in top speed mode with 2 s cycles.

The raw data acquired was analysed using MaxQuant software version 1.5.51 allowing 10 ppm mass accuracy on precursor masses and 0.8 Da on fragment ions. Tryptic petpides were specificed with two missed cleavages and a mixed modification of carbamidoethylation on Cys residues and variable oxidation on Met and acetylation on Lysine residues. The UNIPROT Drosophia melanogaster databse from August 2017 was searched containing 22113 entries and a additional database of common contaminating proteins (keratin, trypsin) was also searched. Scaffold (TM, version 4.6.2, Proteome Software Inc.) was used to validate MS/MS based peptide and protein identifications. Peptide identifications were accepted if they could be established at greater than 95.0% probability by the Scaffold Local FDR algorithm. Protein identifications were accepted if they could be established at greater than 99.0% probability and contained at least 2 identified peptides.

### ChIP assay

Approximately 200 whole larvae from wild-type (WT) flies were fixed with 1% Formaldehyde (Sigma-Aldrich, Cat#: F8775) at 37°C for 15 min followed by incubation on ice for 2 min. For the supernatant preparation 200 µl of lysis buffer was added (50 mM Tris-HCl, pH 7.6, 1 mM CaCl_2_, 0.2% Triton X-100 or NP-40, 5 mM butyrate, and 1X proteinase inhibitor cocktail). The lysates were subjected to sonication to shear DNA to the length of approximately between 150 and 900 bp using an Epishear^TM^ Probe Sonicator (Active motif). The sample was then diluted by adding RIPA buffer (10 mM Tris-HCl, pH 7.6, 1 mM EDTA, 0.1% SDS, 0.1% Na-Deoxycholate, 1% Triton X-100, supplemented with protease inhibitors and PMSF). For the input samples 40 µl were saved and supplemented with 2 µl 5M NaCl and were incubated at 65°C O/N to reverse crosslink. The rest of the lysate was then incubated with control IgG (Cell Signaling Technology, Cat#: 2729S) or primary antibody against Sequoia (gift from Prof Y Jan) together with 40 µl of Protein A beads (GE Healthcare, 28967062) at 4ºC O/N. The beads were washed sequentially with the following buffers: RIPA buffer, RIPA buffer supplemented with 0.3M NaCl, LiCl buffer (0.25 M LiCl, 0.5% NP40, 0.5% NaDOC), TE buffer supplemented with 0.2% Triton X-100 and TE buffer. To reverse crosslink, the beads were resuspended in 100 µl TE buffer (supplemented with 10% SDS and Proteinase K (20 mg/ml) and were incubated at 65°C O/N. The DNA was purified by phenol-chloroform extraction followed by qPCR analysis. Primers are listed in Table S2.

### Structural modelling

The model of the peptide of Sequoia binding to *Drosophila* ATG8a was derived from the structure of GABARAP-L1 ATG4B LIR Complex (PDB code 5LXI). Essentially all residues in the structure were mutated to the correct sequence. No energy minimization was carried out. The structures and electrostatic surfaces were displayed in PyMol. As an approximation the electrostatic surface associated with the acetylated protein was calculated with a methionine as a mimic of the acetylated K46.

## Supplementary Figure Legends

**Supplementary Figure 1:**
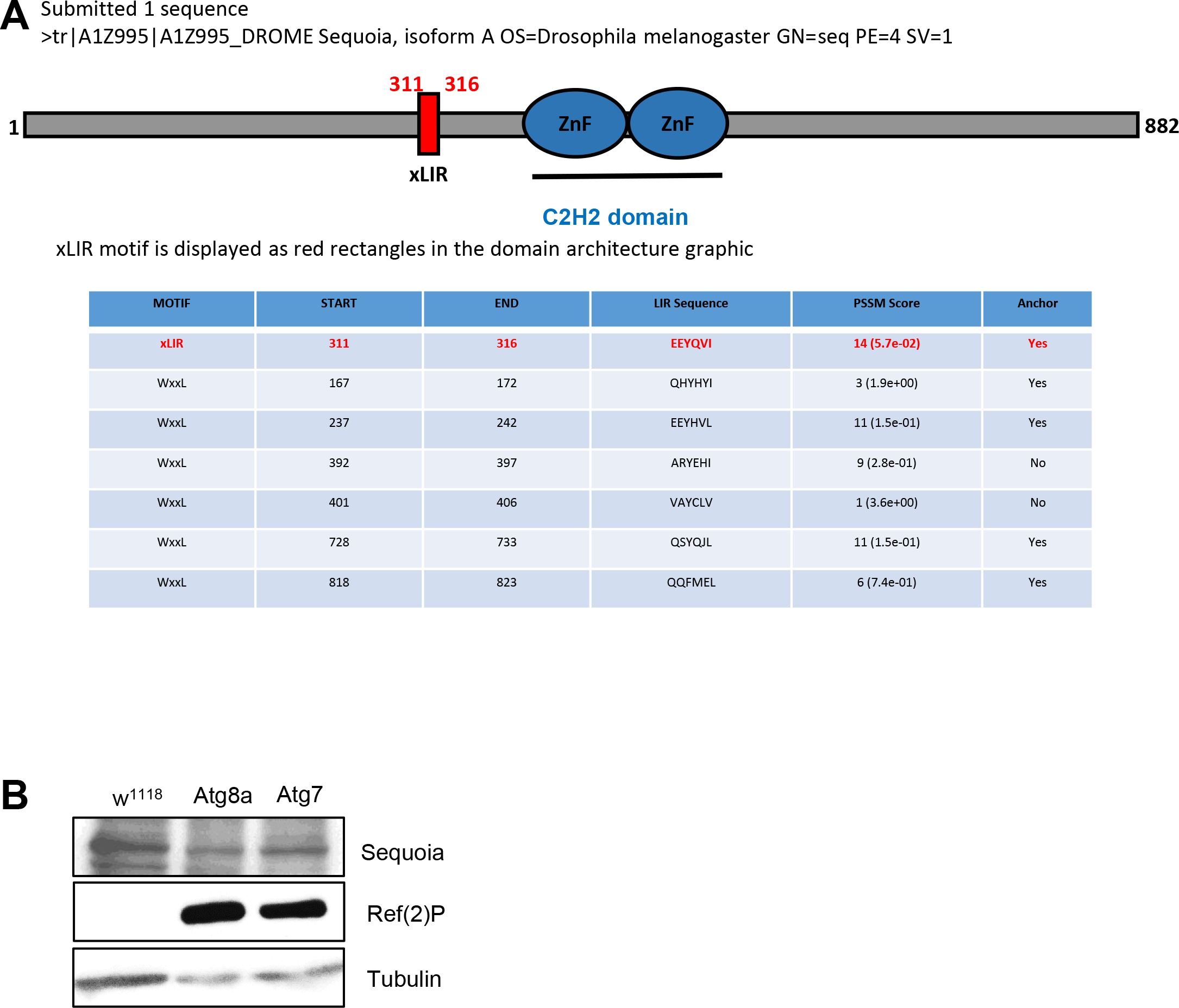
(A) Sequoia is a Zinc finger domain-containing protein with a xLIR motif. Screenshot of the result window from the iLIR server (http://ilir.warwick.ac.uk/) for the protein sequence of Sequoia (Uniprot A1Z995). xLIR relaxed LIR motif; WxxL: conventional LIR motif; ZnF: Zinc finger motif; PSSM: position-specific scoring matrix. (B) Whole body lysates from wild-type (WT) and Atg8a and Atg7 mutant flies were subjected to SDS-PAGE and immunoblotting for Sequoia and Ref(2)P. Tubulin was used as loading control.

**Supplementary Figure 2:**
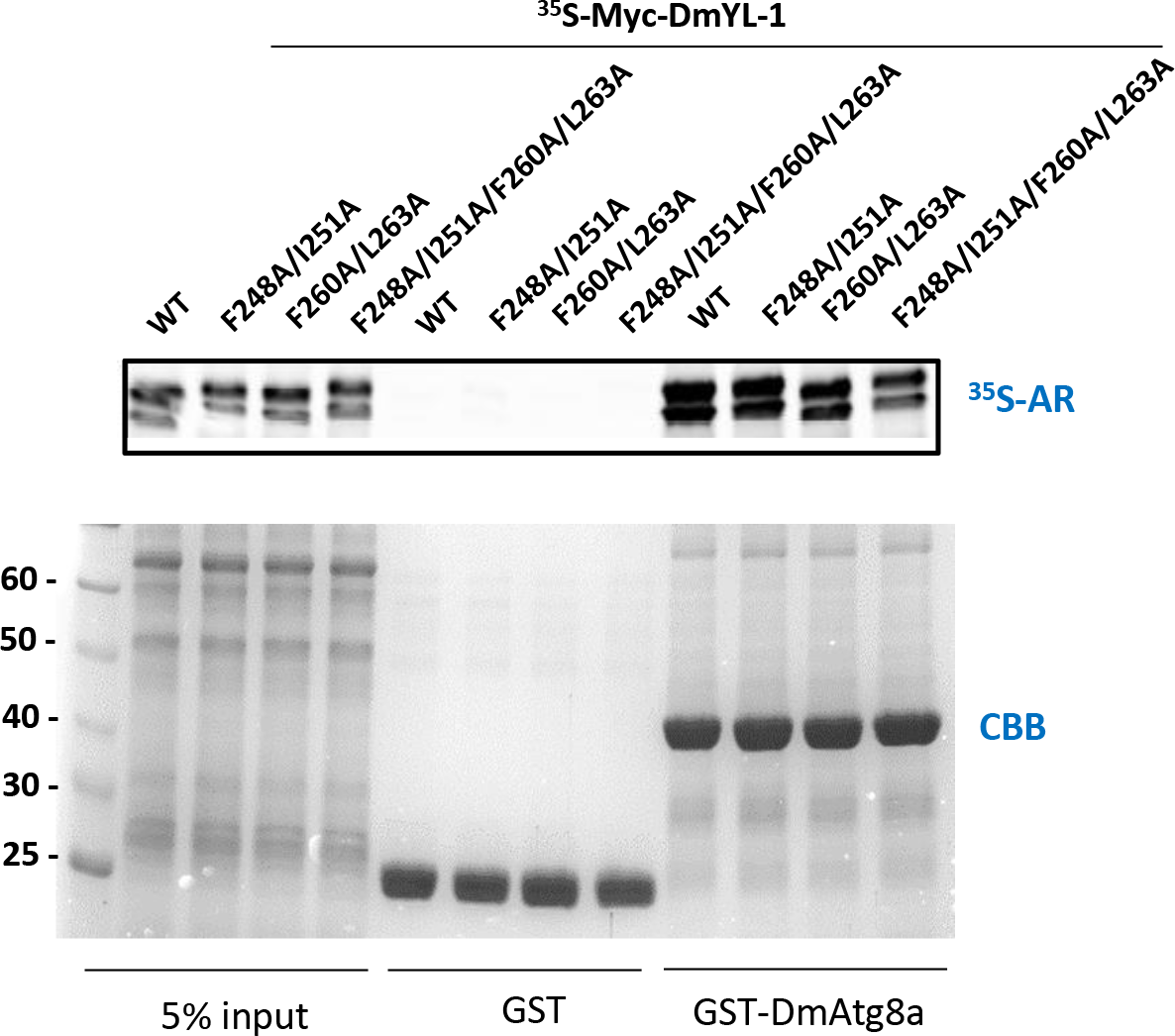
GST-pull-down assay between GST-tagged Atg8a-WT, and putative LIR-mutant constructs and radiolabelled myc-YL-1 WT. AR: autoradiography, CBB: Coommassie blue staining.

**Supplementary Figure 3:**
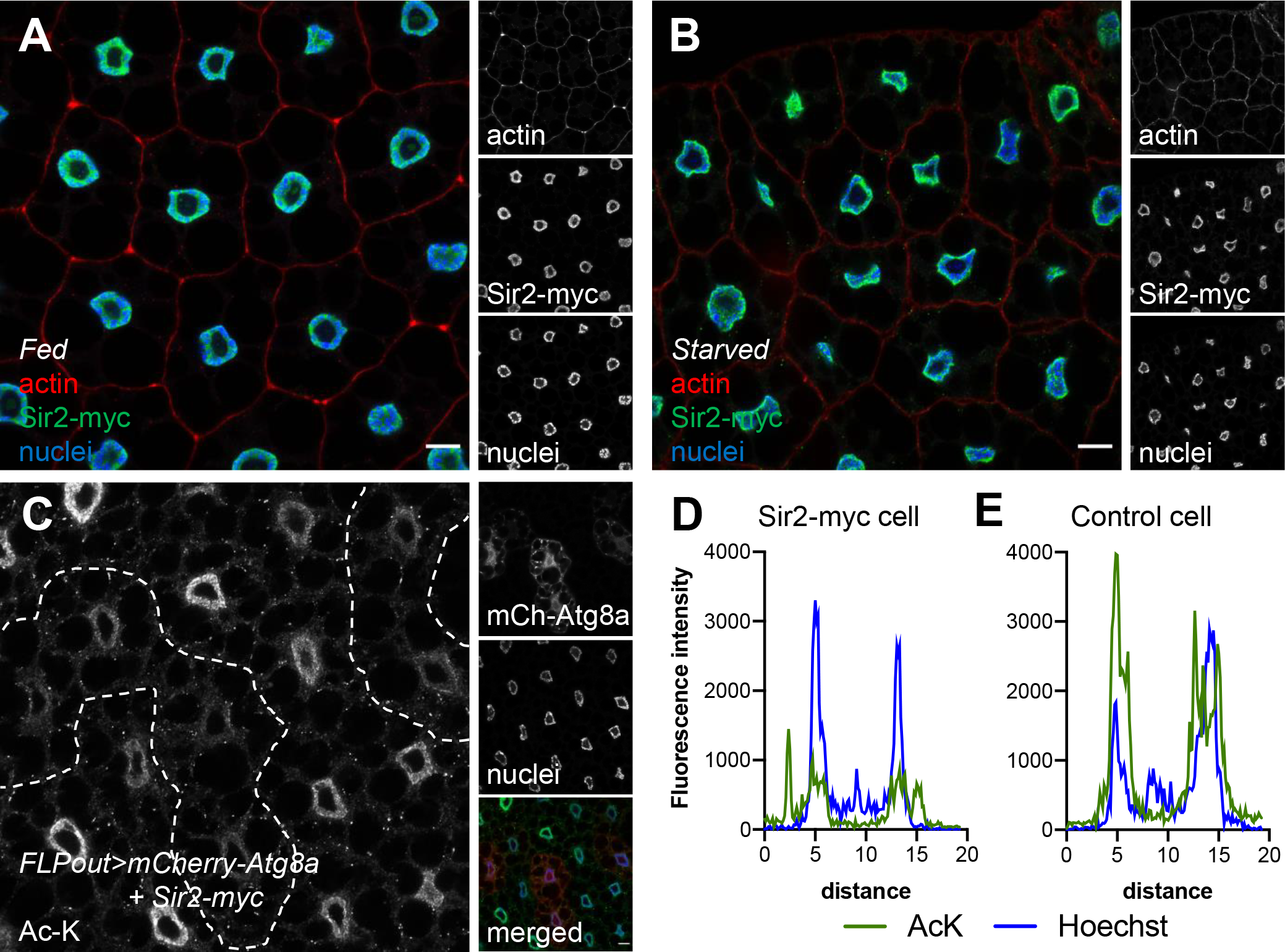
Sir2 deacetylates nuclear proteins. (A-B) Confocal sections of fat bodies expressing Sir2-myc (green) from fed (A) and 4hrs starved (B) larvae. Fixed fat bodies where stained nuclei (blue) and cortical actin (red). (C) Confocal sections of larval fat bodies clonally expressing Sir2-myc in combination with the autophagy marker mCherry-Atg8a (red) and stained for acetylated-lysine (green). (D-E) Intensity plots for the green (acetylated-lysine, AcK) and blue (Hoechst) channels taken across nuclei from control (D) or Sir2-myc clonal (E) cells.

