## Supplementary material for "Atg8a interacts with transcription factor Sequoia to control the expression of autophagy genes in *Drosophila*": Supllementary figures

### Supplementary Figure Legends:

#### Supplementary Figure 1:

(A) Sequoia is a Zinc finger domain-containing protein with a xLIR motif. Screenshot of the result window from the iLIR server (http://ilir.warwick.ac.uk/) for the protein sequence of Sequoia (Uniprot A1Z995). xLIR relaxed LIR motif; WxxL: conventional LIR motif; ZnF: Zinc finger motif; PSSM: position-specific scoring matrix. (B) Whole body lysates from wild-type (WT) and Atg8a and Atg7 mutant flies were subjected to SDS-PAGE and immunoblotting for Sequoia and Ref(2)P. Tubulin was used as loading control.

#### Supplementary Figure 2:

GST-pull-down assay between GST-tagged Atg8a-WT, and putative LIR-mutant constructs and radiolabelled myc-YL-1 WT. AR: autoradiography, CBB: Coommassie blue staining.

#### Supplementary Figure 3:

Sir2 deacetylates nuclear proteins.

(A-B) Confocal sections of fat bodies expressing Sir2-myc (green) from fed (A) and 4hrs starved (B) larvae. Fixed fat bodies where stained nuclei (blue) and cortical actin (red).

(C) Confocal sections of larval fat bodies clonally expressing Sir2-myc in combination with the autophagy marker mCherry-Atg8a (red) and stained for acetylated-lysine (green).

(D-E) Intensity plots for the green (acetylated-lysine, AcK) and blue (Hoechst) channels taken across nuclei from control (D) or Sir2-myc clonal (E) cells.

**A** Submitted 1 sequence
>tr|A1Z995|A1Z995_DROME Sequoia, isoform A OS=Drosophila melanogaster GN=seq PE=4 SV=1

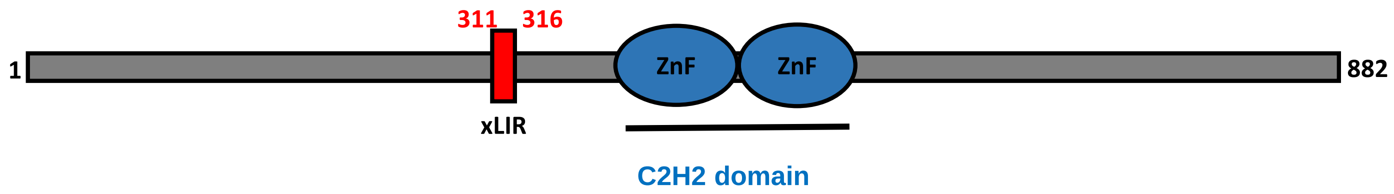


xLIR motif is displayed as red rectangles in the domain architecture graphic

| MOTIF | START | END | LIR Sequence | PSSM Score | Anchor |
|---|---|---|---|---|---|
| xLIR | 311 | 316 | EEYQVI | 14 (5.7e-02) | Yes |
| WxxL | 167 | 172 | QHYHYI | 3 (1.9e+00) | Yes |
| WxxL | 237 | 242 | EEYHVL | 11 (1.5e-01) | Yes |
| WxxL | 392 | 397 | ARYEHI | 9 (2.8e-01) | No |
| WxxL | 401 | 406 | VAYCLV | 1 (3.6e+00) | No |
| WxxL | 728 | 733 | QSYQJL | 11 (1.5e-01) | Yes |
| WxxL | 818 | 823 | QQFMEL | 6 (7.4e-01) | Yes |

**B**

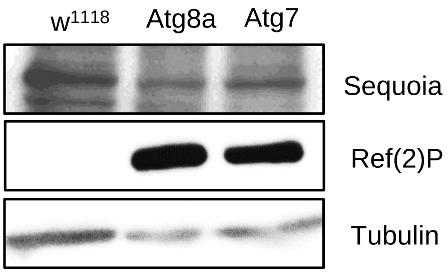

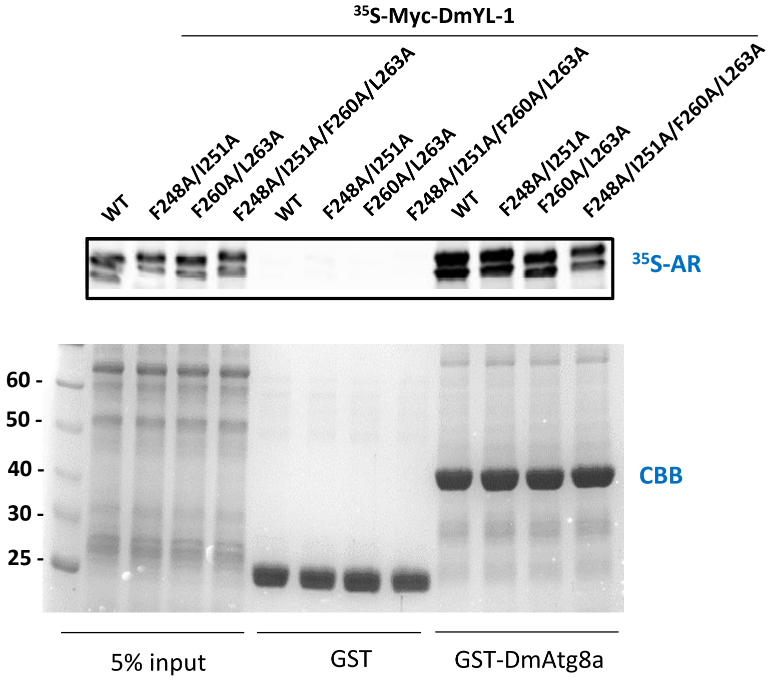
35S-Myc-DmYL-1
WT
F248A/I251A
F260A/L263A
F248A/I251A/F260A/L263A
WT
F248A/I251A
F260A/L263A
F248A/I251A/F260A/L263A
WT
F248A/I251A
F260A/L263A
F248A/I251A/F260A/L263A
35S-AR
60 -
50 -
40 -
30 -
25 -
CBB
5% input
GST
GST-DmAtg8a

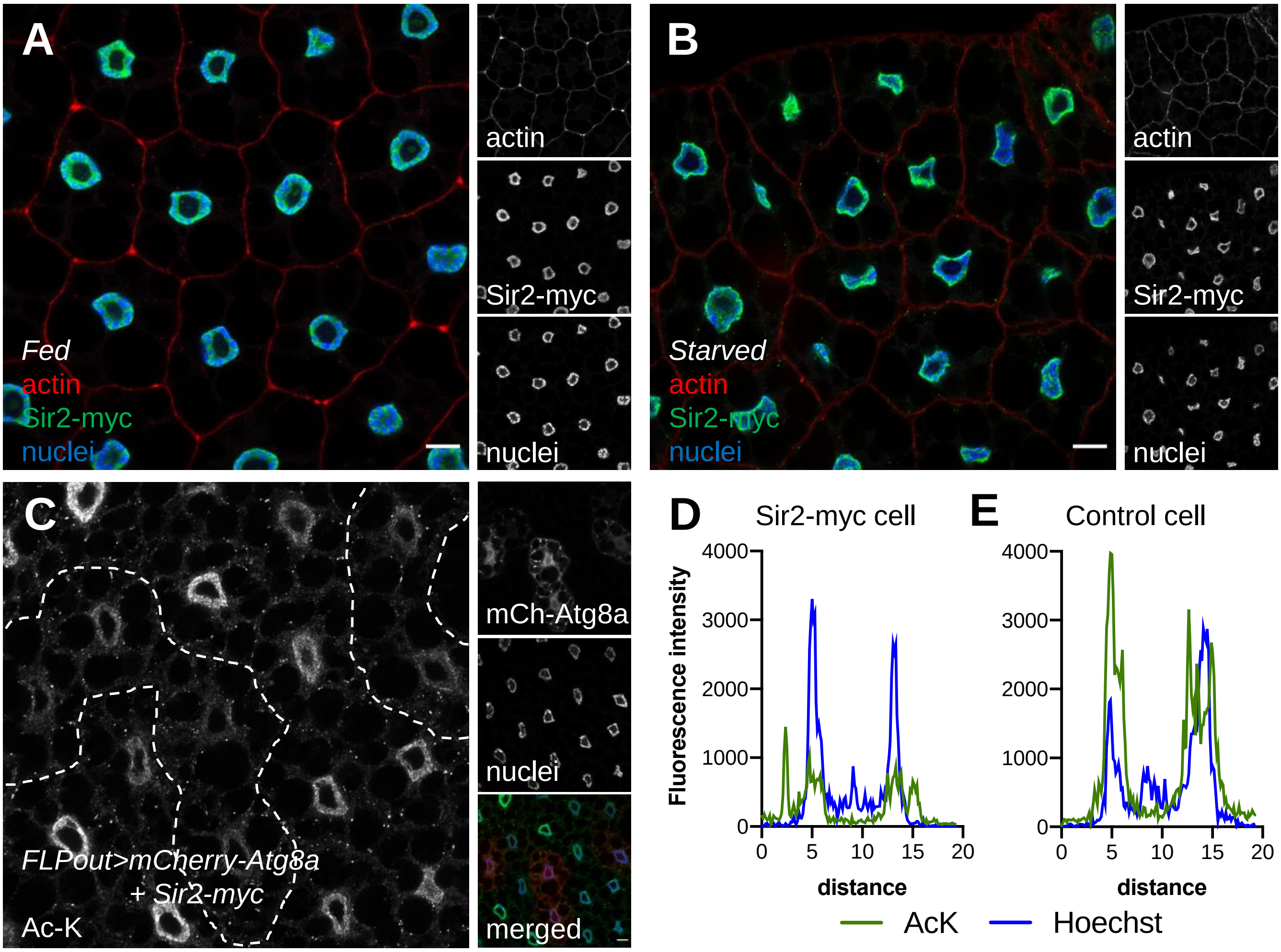

A
Fed
actin
Sir2-myc
nuclei
actin
Sir2-myc
nuclei
B
Starved
actin
Sir2-myc
nuclei
actin
Sir2-myc
nuclei
C
FLPout>mCherry-Atg8a
+ Sir2-myc
Ac-K
mCh-Atg8a
nuclei
merged
D
Sir2-myc cell
E
Control cell
Fluorescence intensity
4000
3000
2000
1000
0
0 5 10 15 20
distance
AcK
Hoechst
